# Genetic diversity within a tree and alternative indexes for different evolutionary effects

**DOI:** 10.1101/2024.02.24.581556

**Authors:** Yoh Iwasa, Sou Tomimoto, Akiko Satake

## Abstract

Trees, living for centuries, accumulate somatic mutations in their growing trunks and branches, causing genetic divergence within a single tree. Stem cell lineages in a shoot apical meristem accumulate mutations independently and diverge from each other. In plants, somatic mutations can alter the genetic composition of reproductive organs and gametes, impacting future generations. To evaluate the genetic variation among a tree’s reproductive organs, we consider three indexes: mean pairwise phylogenetic distance 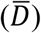, phylogenetic diversity (*PD*; sum of branch lengths in molecular phylogeny), and parent-offspring phylogenetic distance (*D*_*PO*_). The tissue architecture of trees facilitated the accumulation of somatic mutations, which have various evolutionary effects, including enhancing fitness under strong sib competition and intense host-pathogen interactions, efficiently eliminating deleterious mutations through epistasis, and increasing genetic variance in the population. Choosing appropriate indexes for the genetic diversity of somatic mutations depends on the specific aspect of evolutionary influence being assessed.

## 1. Introduction

Trees have long lifespans ranging from tens to hundreds of years, during which their trunks and branches continue to grow. Somatic mutations occur due to errors in genome replication during cell division and failures in DNA damage repair following physical or chemical disturbances. These mutations arise and accumulate in different branches, leading to genetic differentiation within an individual tree (Tomimoto & Satake 2023). Thanks to recent developments in genomic analysis technologies such as next-generation sequencing, we have become able to identify genetic patterns caused by somatic mutations (Schmidt-Siegert et al. 2017; Plominon et al. 2018; Hanlon et al. 2019; Wang et al. 2019; Orr et al. 2020; Hofmeister et al., 2020; Zahrdadníková et al. 2020; Reusch et al. 2021; Perez-Roman et al., 2022; Duan et al. 2022; Schmitt et al. 2022). The physical structure of the tree, with its branches, exhibited a topology similar to the molecular phylogenetic trees of cells sampled from different branches (Orr et al., 2020; Satake et al. 2023), suggesting that somatic mutations accumulate as shoots elongate.

In plant tissues, somatic genetic variations can contribute to the next generation because reproductive organs (flowers and fruits) originate from stem cells in the shoot apical meristems (abbreviated as SAM). Gametes, including eggs and pollen, can undergo genetic diversification through somatic mutations, thereby contributing to genetic variation among offspring (Whitham & Slobodchikoff 1981; Sutherland & Watkins 1986; Plomion et al. 2018). This differs from animals with unitary structures, where germ lines differentiate from soma in early developmental stages.

In a previous paper (Iwasa et al. 2023), we studied a mathematical model describing how the genetic pattern of a shoot is determined by the behavior of stem cells in the meristem. We evaluated phylogenetic distance between cells sampled from different portion of a shoot, indicating their genetic difference due to somatic mutations accumulated during shoot elongation. Stem cells in the SAM normally undergo asymmetric cell division, producing successor stem cells and differentiated cells. However, occasionally, a stem cell may fail to leave its successor stem cell. Subsequently, to recover stem cell number, the vacancy is filled by the duplication of one of the nearest neighbor stem cells. Because cell walls prevent stem cells from exchanging their positions, this leads to the genetic differentiation of cells according to the angle around the shoot and a larger genetic variance accumulated among cells in the body (Iwasa et al. 2023).

### 1.1 Characteristics of tissue structure of trees

Here, we highlight several characteristics of tree tissue structure, some of which are shared with modular animals (e.g., corals and sponges; Vasquez Kuntz et al. 2022).

i. No clear distinction between germ line and soma: In plant tissues, somatic mutations accumulated during shoot elongation may contribute to reproductive organs and gametes (eggs and pollen), affecting the genetic content of offspring.
ii. Shoot apical meristems on different branches: In trees, each shoot (trunk or branch) contains a shoot apical meristem (SAM) with a small number of stem cells. The SAMs are physically separated and located at the tips of different branches. Stem cells in different SAMs remain isolated. This structure facilitates independent accumulation of somatic mutations between branches, leading to their genetic differentiation.
iii. Before-forking portion of ancestral cell lineages: Stem cells undergo asymmetric cell division, creating both a successor stem cell and a differentiated cell. The differentiated cells then undergo a finite number of duplications, increasing in number and size to form a portion of a shoot. If the stem cells perform asymmetric cell division with a high probability, different stem cell lineages accumulate somatic mutations independently, and stem cells in the SAM become genetically diverse. The ancestral lineages of two different stem cells in the SAM have some distance between their common ancestor and the sampled location as illustrated in Fig. 1. This is referred to as “before-forking” portion (Tomimoto et al. 2023). Its magnitude depends on the probability of failure in asymmetric cell division in stem cells (Iwasa et al. 2023).

**Fig. 1.**
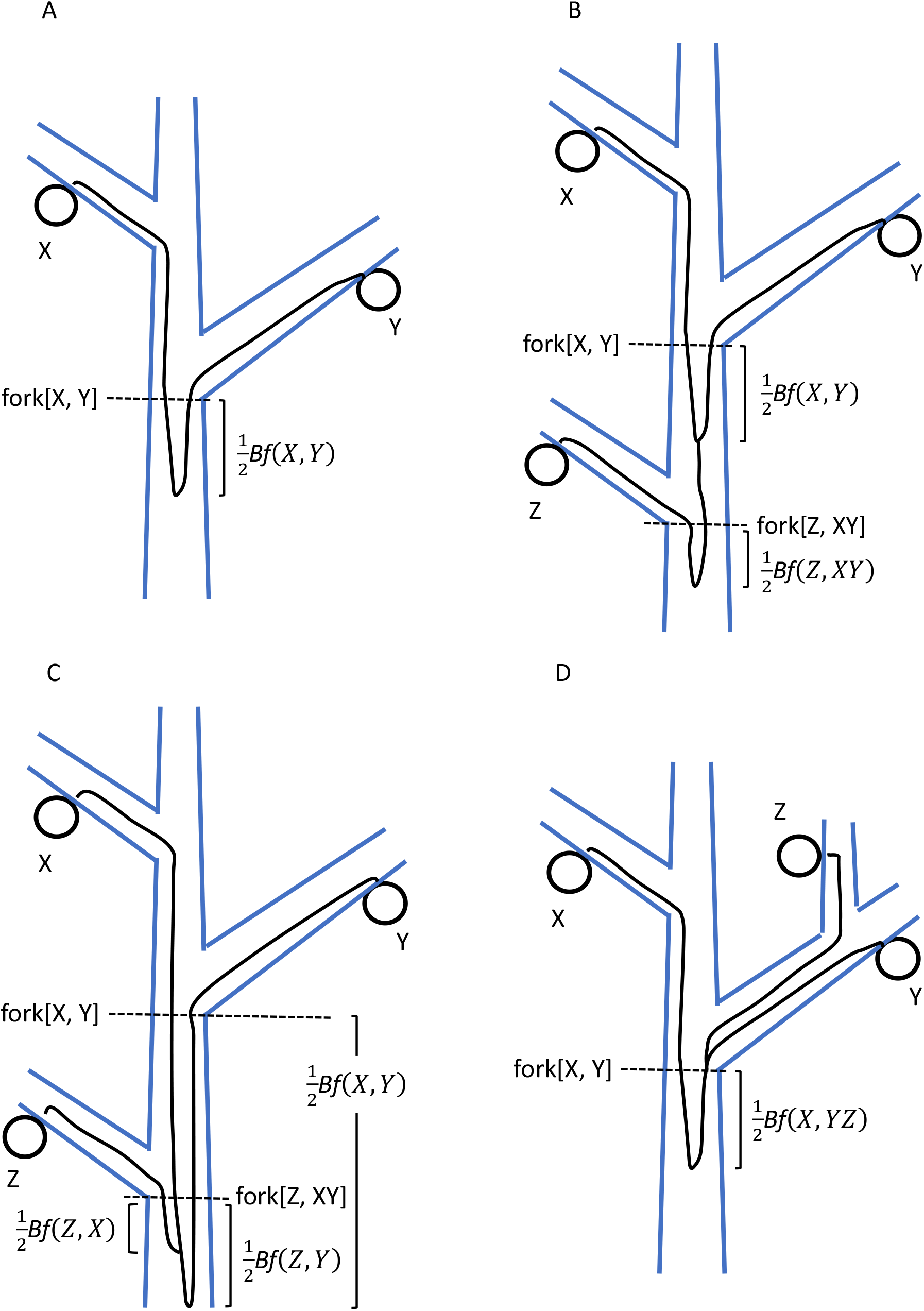
Scheme of ancestral cell lineages connecting sampled cells on different branches of a tree. Notation of symbols and explanations are provided in the main text and Part A of the SM. (A) X and Y are sampled cells. The ancestral cell lineages of X and Y converge at the same location, marked as fork[X, Y]. Their ancestral cell lineages then merge below it at a distance of 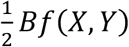. (B) Three cells, X, Y, and Z, are sampled from different branches. The coalescence of ancestral lineages of X and Y occurs below fork[X, Y] at a distance of *Bf*(*X, Y*). It takes place above fork[Z, XY], which represents the forking between ancestral lineage of Z and the common ancestral cell lineage of X and Y. The coalescence of Z and XY occurs below fork[Z, XY] at a distance of 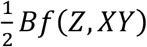. (C) Three cells X, Y, Z are sampled. The coalescence of A and B does not occur above fork[Z, XY], where the shoot apocal meristem (SAM) includes three ancestral cells of X, Y, and Z. (D) The coalescence of Y and Z does not occur within the lateral branch. Instead, it occurs at fork[X, YZ] as a result of bottleneck in the stem cell population in the SAM when the lateral branch is spliced.
iv. Circular genetic differentiation around a shoot: When a stem cell fails to leave its successor, the stem cell line is replaced by a copy of a neighboring stem cell, as cell movement is constrained by the cell wall. This produces circular genetic differentiation of stem cells and leads to larger genetic diversity among them than when they are well mixed (Iwasa et al. 2023). Possible roles of the layer structure in the meristem have also been discussed (Klekowski & Kazarinova-Fukshansky 1984; Klekowski et al. 1985; Pineda-Krch & Lehtilä 2002).
v. Distinction between the main and lateral branches: When a lateral branch is formed at a main branch, the stem cells of the axillary meristem of the lateral branch are sampled from the SAM of the main branch. In many trees, only a small fraction of stem cells in the main branch contribute to the axillary meristem (Tomimoto & Satake 2023; Tomimoto et al. 2023). This results in an asymmetry between the two stem cell lineages sampled above the forking. As illustrated in Fig. 1D, the main branch maintains multiple ancestral cell lineages just below the forking, while cell lineages from the lateral branch coalesce at the forking. We may be able to distinguish the main and lateral branches at a forking by comparing the genetic diversity among cells of the two branches.

In the present paper, as a sequel to Iwasa et al. (2023), we explore the genetic diversity among reproductive organs on different branches of a single tree. We examine three indexes for the magnitude of the diversity: the mean pairwise phylogenetic distance, the phylogenetic diversity, and parent-offspring phylogenetic distance. These indexes are calculated from the length of ancestral stem cell lineages between sampled cells. We discuss their suitability for measuring the magnitude of genetic variations caused by somatic mutations, considering their evolutionary effects.

## 2. Genetic distance between cells and the length of ancestral cell lineages

We consider the following situation: A shoot consists of cells derived from a small number of stem cells in the SAM located at the tip. A stem cell undergoes asymmetric cell division and produces one successor stem cell and one differentiated cell. The differentiated cells increase in number through a finite number of cell divisions, grow in cell size, form a portion of the shoot, and lift up the SAM. This process continues and creates a shoot with the current SAM on the tip and the lower portion of the shoot reflecting an earlier genetic state of stem cells in the SAM (Iwasa et al. 2023).

All cells forming above-ground plant body are ultimately derived from stem cells in the SAM. As the shoot elongates, novel mutations accumulate. Hence, the genetic distance between two cells can be evaluated by the length of the ancestral lineage connecting them. Iwasa et al. (2023) defined phylogenetic distance *D* as the physical length along a branch, measured in units of cm or m. The genetic distance, which represents the expected number of genomic differences between cells, is calculated by multiplying *D* by the mutation rate per physical shoot length. Mutations occur due to errors in cell division and failures of DNA damage repair, with the rates proportional to the number of cell divisions and to the calendar time, respectively. Comparing trees between fast and slow growing tropical species, Satake et al. (2023) concluded that the mutation rate proportional to the number of calendar years (independent of the number of cell divisions) was much more important than the other type. While Iwasa et al. (2023) discussed *D* between two cells sampled from the same shoot, in the present paper, we consider the cases where the sampled cells are on different branches of a tree.

Fig. 1A illustrates an example of ancestral cell lineages connecting two sampled cells X and Y, located on different branches that are spliced from the main branch (trunk). The location labeled as fork[X, Y] represents the forking point between X and Y, where their ancestral cell lineages of cells X and Y converged to the same SAM. These two ancestral stem cells may or may not be identical, because a SAM contains multiple stem cells. If they are different, there is a distance between their common ancestor stem cell and fork[X, Y]. We call this portion as “before forking” segment (Tomimoto et al. 2023) and denote the length of the path between two ancestral stem cells at fork[X, Y] by *Bf*(*X, Y*). The physical distance between the common ancestor cell and fork[X, Y] is half of *Bf*(*X, Y*) and was called “coalescent length” between the two ancestral cells at fork[X, Y] (Iwasa et al. 2023)

The before-forking portion may extend to the base of the shoot. However, there is a possibility that their common ancestral stem cell exists in the middle of the shoot. This phenomenon, known as “coalescence”, can occur because a stem cell in the SAM sometimes fails to leave its successor stem cell. When such a failure occurs, another stem cell in the SAM duplicates to replenish the stem cell count. Consequently, a stem cell line is replaced by another. As a result, the ancestral cell lineage of a focal stem cell shifts its location (Iwasa et al. 2023).

The phylogenetic distance between X and Y is defined as the length of their stem cell lineage since their common ancestor cell (Fig. 1A), given by:

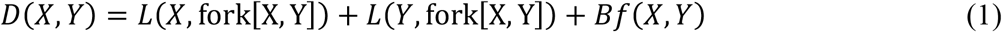

where the first term on the right-hand side represents the physical length along the branch between sampled cell X and the forking (see Iwasa et al., 2023 for definition of

*L*). The second term represents a similar quantity for sampled cell Y. Detailed explanations are provided in Part A of the Supplemental Material (abbreviated as SM). The sum of the first and second terms on the right-hand side of Eq. (1) is the after-forking portion of the phylogenetic distance. The last term indicates the before-forking term, which increases with the genetic diversity of stem cells within the SAM.

Fig. 1B illustrates the case of three sampled cells from different branches. The ancestral cell lineages of X and Y coalesce below fork[X, Y] by 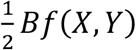. Their common ancestral cell lineage then continues downward. The forking between their common ancestral lineage and the ancestral cell lineage of Z is denoted by fork[Z, XY]. In Fig. 1B, the common ancestor of X and Y is above fork[Z, XY]. Coalescence of all three ancestral lineages occurs where the ancestral lineage of Z and the common ancestral lineage of X and Y coalesce. It is located below fork[Z, XY] by length 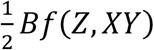. For further details, refer to Part A of the SM.

Fig. 1C illustrates the case ancestral cell lineages of X and Y do not coalesce above fork[Z, XY]. The ancestral cell lineages have three different ancestral cells at fork[Z, XY]. We observe a coalescent process of these three cells. In Fig. 1C, the ancestral cell lineages of X and Z coalesce first, followed by the coalescence of their common ancestral lineage with the ancestral lineage of Y.

Fig. 1D illustrates the asymmetry between main and lateral branches. The ancestral cell lineages of Y and Z do not coalesce within the lateral branch but they coalesce where the lateral branch is spliced, because the stem cells of the SAM of the lateral branch were assumed to derive from a single stem cell in the SAM of the main shoot. In contrast, the main branch keeps the diversity, as illustrated by that the ancestral cell lineage of X was kept separate from the common ancestral cell lineage of Y and Z.

### 2.1 Mean pairwise phylogenetic distance between reproductive organs 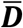

A tree may have many reproductive organs, producing seeds or flowers, with the latter producing pollen to fertilize the seeds of other trees. Let *n* be the number of reproductive organs of a tree, which may vary in the number of offspring they produce. We represent relative sizes of these organs as ω_1_, ω_2_, . . ., ω_*n*_, which satisfy ω_1_ + ω_2_+. . . +ω_*n*_ = 1. The phylogenetic distance between organs *i* and *j* is denoted as *D*(*i, j*), which is defined by Eq. (1) (see Fig. 3A).

Here, we define the mean pairwise phylogenetic distance among all the reproductive organs of an individual tree as follows:

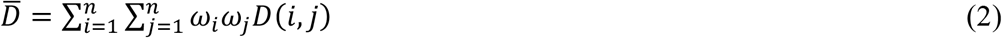

This equals to the mean phylogenetic distance between two randomly chosen seeds.

The genetic diversity is quantified by 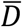 given in Eq. (2), multiplied by the mutation rate per unit shoot length.

Cells from the focal tree undergo meiosis, become haploid gametes, experience syngamy with the gametes produced by other trees, form seeds, and become young trees in the next generation. Hence, tree structure with a larger 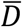 realizes genetically more diverse offspring.

## 3. Phylogenetic diversity of multiple stem cells of a tree *PD*

Phylogenetic diversity (abbrev. *PD*) is defined as the sum of branch lengths of phylogeny (Fig. 3B). It was first proposed by Faith (1992) as a measure of the importance of different species (or taxonomic units) in conservation biology, and has been adopted widely to evaluate the diversity of a group of multiple species (microbes; Sogin et al., 2006; Lauber et al. 2009). In the classical gene-genealogy theory, the sum of branch lengths of molecular phylogeny is calculated for the case that genomes are sampled from a single well mixed population. Watterson (1975) defined the expected number of segregating sites, denoted by *S*_*n*_, as the product of mutation rate *m* and the sum of branch lengths in gene phylogeny.

We apply *PD* to quantify the genetic diversity of reproductive organs of a single individual tree, treating them as taxonomic units. This approach can be seen as an adaptation of Waterson’s total length of molecular phylogeny to the population of stem cells. However, it is important to note that sampling was conducted in a population with geographic structures characterized by separation into multiple SAMs at different branches and a circular genetic structure within each SAM.

Both mean pairwise phylogenetic distance 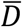 and phylogenetic diversity *PD* measure the genetic differences between reproductive organs and evaluate the variability among offspring (i.e., seeds and young trees). However, these two metrics emphasize different aspects of the genetic diversity among offspring. The phylogenetic diversity *PD* is proportional to the total number of haplotypes included in the whole phylogeny, giving equal weight to genotypes that produce numerous seeds and those that produce a single seed. In contrast, the mean phylogenetic distance between organs 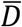 considers branches with many seeds more important than those with few sees.

Additionally, even when all the reproductive organs have equal weight (ω_1_ = ω_2_ =. . . = ω_*n*_ = 1*/n*), 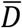 and *PD* assign different weights to various branches. Fig. 2 illustrates a tree with eight reproductive organs (*n* = 8). In Fig. 2A, each branch has an importance proportional to the number of ancestral lineages connected two organs. Note that removing a branch results in two disconnected trees. If the number of organs included in these trees are *i* and *j*, the weight for the branch is proportional to *ij*. In Fig. 2A, we indicated the importance of each branch in calculating 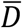, with the values normalized to ensure their sum equals 100. In contrast, Fig. 2B illustrates the equal importance of each branch for *PD*, with the same value assigned to all branches. Comparing these two, we can conclude that 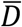 assigns more weight to branches in the central portion than those in the marginal portion of the tree (molecular phylogeny). For more detailed explanations of *PD*, refer to Part B of the SM.

**Fig. 2.**
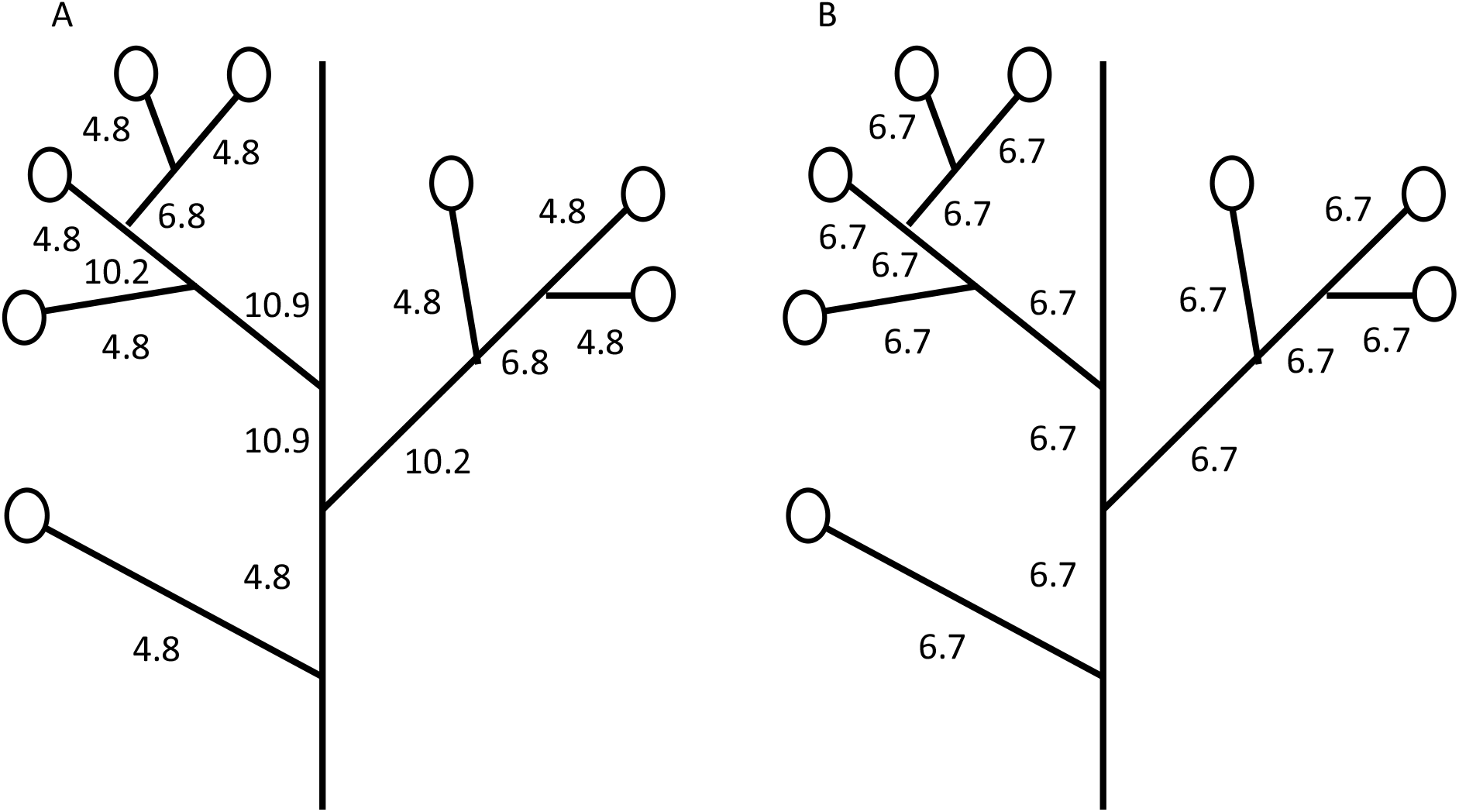
Difference in the relative importance of branches in a tree when calculating 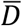 and *PD*. Open circles indicate reproductive organs (flowers and fruits) on the branches of a tree. In this scenario, the same number of seeds is produced per reproductive organ, and their weights are equal: ω_1_ =. . . = ω_8_ = 0.125. (A) The relative importance of branches in calculating 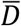. They are normalized to ensure their sum equals 100. In this case, the importance of a branch is proportional to the number of ancestral cell lineages connecting two reproductive organs. (B) The relative importance of branches in calculating *PD*.

**Fig. 3.**
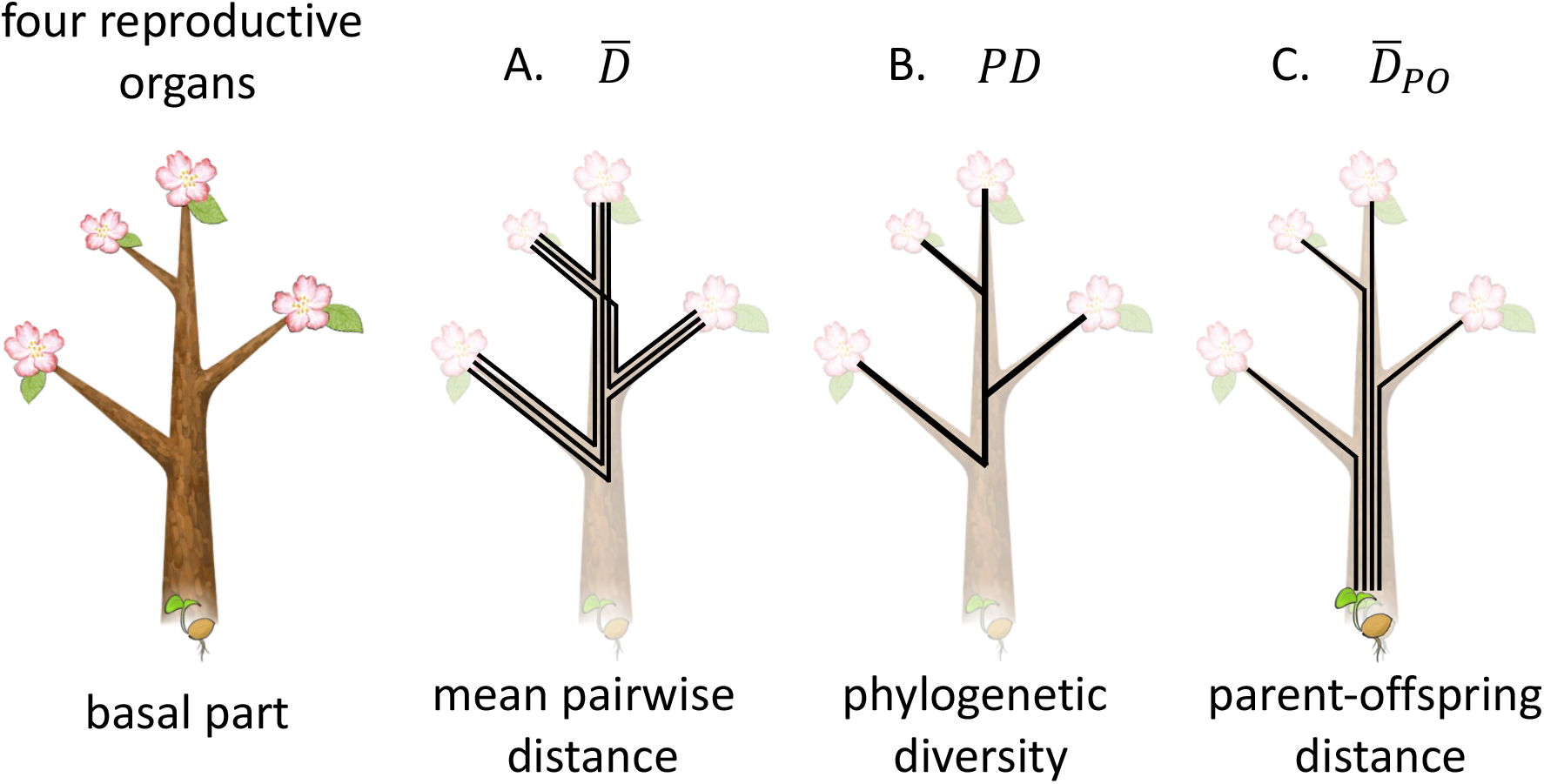
Three indexes for the genetic diversity of multiple reproductive organs of a single tree. We consider a tree with four reproductive organs on different branches (see the figure on the left). The three different indexes are: A. mean pairwise phylogenetic difference 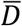 ; B. Phylogenetic diversity *PD*; and C. parent-offspring genetic distance *D*_*PO*_. Refer to the text for the explanations.

### 3.1 Phylogenetic diversity of stem cells sampled at the same position

Fig. 1C illustrates the case where the ancestral cells of three sampled cells, X, Y, and Z, are different cells in the same SAM. We need to evaluate the sum of branch lengths of ancestral lineages of these three cells until the coalescence. In this section, we consider the phylogenetic diversity of more than two stem cells in the same SAM along the shoot.

In Part B of the SM, we discuss the mean value of phylogenic diversity of sampled stem cells. Let *n* be the number of stem cells in the SAM. They are arranged in a circular manner, with the nearest neighbors separated by an angle of 2π*/n* radians. The number of sampled stem cells is *m*, and they are separated with their neighbors by *k*_1_, *k*_2_, . . ., *k*_*m*_ (*k*_1_+. . . +*k*_*m*_ = *n*). If a stem cell failed to leave its successor stem cell, the location is filled by a copy of its nearest neighbor. This occurs at a rate *q* per unit shoot length. Then the location of the ancestral lineage of a sampled cell shifts by one, either to the left or right. The mean phylogenic diversity of sampled stem cells, denoted by *PD*(*k*_1_, . . ., *k*_*m*_; *y*), depends on *y*, the distance of the sampled location from the base of the shoot.

In Part B of the SM, we derive a differential equation for *PD*(*k*_1_, . . ., *k*_*m*_; *y*) and obtain the solution considering the boundary conditions. Fig. 4 illustrates how *PD*(*k*_1_, . . ., *k*_3_; *y*) increases with *y*. In the limit of very large *y* (*y* → ∞), we have the following solution:

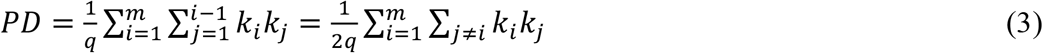

which indicates the total sum of the coalescent lengths of *m* stem cells if the sampled location is far from the base of the shoot. *PD* is large when stem cells have a high probability of leaving their successor cells (*q* is small). It also depends on the intervals between the sampled stem cells (*k*_1_, . . ., *k*_*m*_). We can rewrite Eq. (3) as follows:

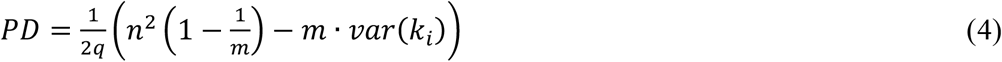

where *var*(*k*_*i*_) is the variance of the length of intervals: Phylogenetic diversity reaches its maximum when the sampled cells are equally spaced. When the stem cells are sampled with equal intervals (*var*(*k*_*i*_) = 0), 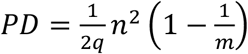 holds.

When we sample just two cells (*m* = 2), *PD* is the same as the phylogenetic distance *D* = *k*(*n* − *k*)*/q*, which was the basis of index 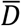 calculated in the last section.

## 4. Parent-offspring distances *D*_*PO*_

Both the pairwise phylogenetic distance 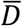 and phylogenetic diversity *PD* are concerning the genetic differentiation among reproductive organs of a single tree, which represent the genetic variation among offspring. The genetic diversity generated by somatic mutations per generation contributes the genetic variation of the whole population, which controls the speed of adaptation of traits under anthropogenetic environmental changes.

The genetic variance of a quantitative trait in the entire population increases by mutations and decreases due to random genetic drift and stabilizing selection (Lande 1976, 1981). The rate at which genetic variance is produced per generation is calculated based on the trait change from the parent to offspring, rather than the difference between offspring. Hence, to evaluate the contribution to the genetic variation of the population, we need to know the phylogenetic distance between parent and offspring (Fig. 3C).

**Fig. 4.**
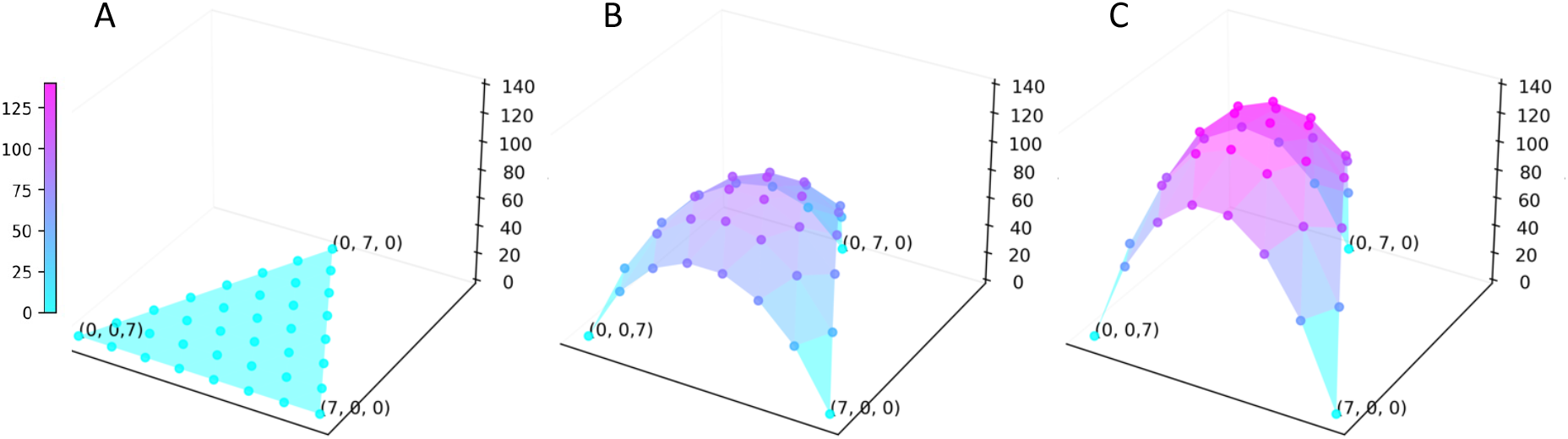
Phylogenetic diversity of three stem cells sampled from the same SAM. The intervals between sampled stem cells are denoted as: *k*_1_, *k*_2_, and *k*_3_, which are positive integers satisfying *k*_1_ + *k*_2_ + *k*_3_ = *n*. Two horizontal axes represent *k*_1_ and *k*_2_. The calculation is made based on the system of differential equations given in Eq. (B.2) in Part B of the SM, showing *PD* = *D*(*k*_1_, *k*_2_, *k*_3_; *y*). Three parts correspond to different distances from the base of the shoot: (A) *y* = 0; (B) *y* = 50; (C) *y* = ∞. Parameters are: *q* = 0.1, *n* = 7, and *m* = 3.

We here consider the parent-offspring phylogenetic distance as follows:

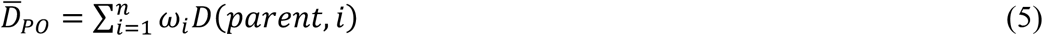

where *D*(*parent, i*) is the phylogenetic distance between the parent genome and the *i*th reproductive organ: *D*(*parent, i*) = *L*(base of the whole shoot, *i*), which is simply equal to the physical length between the base of the whole shoot and the reproductive organs. 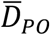 is independent of the genetic diversity of stem cells in the SAM, unlike the two other indexes.

## 5. Discussion

Trees exhibit several characteristic aspects of tissue structure that differ from many animals (unitary organisms). As a result, trees accumulate somatic mutations efficiently, which in addition contribute to the genetic variability of offspring. The selective advantage of somatic mutations expressed in the current generation have been discussed: For example, somatic mutations can lead to diversification in defense chemicals between branches, reducing the harm of herbivores and pathogens (Antolin & Strobeck 1985; Gill et al. 1995; Folse & Roughgarden 2012). In addition, somatic mutations can modify the genome of reproductive organs and the genomic composition of gametes (eggs and pollens), resulting in genetically diverse offspring (Whitham & Slobodchikoff 1981; Schoen & Schultz 2019). These may have diverse evolutionary effects. In this section, we will first address these effects and then discuss how three indexes for the genetic diversity of somatic mutations assess different evolutionary effects.

### 5.1 Generating genetic variations

We may ask whether we can justify the argument that the tissue structure of trees might have evolved as a result of promoting a high rate of mutations in the next generation.

In Part C of the SM, we briefly summarize the conclusions of population genetics theory for the evolution of mutation rates. In the simplest populations (with constant environments over time, no spatial structure, etc.), the mutation rate controlled by neutral modifiers evolves towards lower values (Karlin 1975; Karlin & McGregor 1974; Liberman et al. 2011; Altenberg et al. 2017). In the populations with various structures, a higher mutation rate can evolve. The key elements of these processes were originally studied in the context of the evolution of sex and recombination. Among them, processes such as environmental fluctuations (Ishii et al. 1989; Sasaki & Iwasa 1987), interactions with antagonistic species (Haraguchi & Sasaki 1996, 1997; Sasaki & Haraguchi 2000), spatial heterogeneity, and sibling competition (Williams 1975; Maynard Smith 1978; Bell 1982; Douge & Iwasa 2017a, 2017b) have been shown to potentially favor the evolution a high mutation rate.

Many of these theoretical studies discuss the evolution of the rate of switches between functional alleles that can confer advantages in different environments. However, in most situations, mutations arise due to errors in replication or repair, and they often result in producing many defective and malfunctional genes. Therefore, it is very difficult for a high mutation rate to be advantageous in general circumstances, except for special mechanisms appearing in host-pathogen interactions (Metzger & Willis 2000; Rosenberg et al. 1998). Hence, the observed positive rate of mutation in genomes is considered to have evolved not because of the advantage of higher mutation rate but determined by the high cost of reducing the mutation rate further by investing in more accurate repair or duplication of the genome (Lynch 2010, 2011).

Therefore, we can conclude that the tissue structure of trees has evolved not because a high mutation rate but because other reasons, such as photosynthetic efficiency, competition with neighboring individuals, defense against pathogens or predators, mechanical strength, etc. Mutations produced by trees that contribute to the next generation should be considered as “byproducts” of the tissue structures.

Even if the tissue structure of trees evolved primarily for other reasons, examining the evolutionary effects quantitatively is important. Among many processes that may provide a fitness advantage for having a higher mutation rate, the two most promising ideas are: (1) the antagonistic genotype-specific interaction with pathogens and herbivores, and (2) situations characterized by significant spatio-temporal variation and intense sib-competition (refer to Part C of the SM). These promote the advantage of producing a few diverse offspring, rather than having many similar offspring. It is worthy to quantify these fitness effects in the field. They are likely to be important for genes involved in antagonistic interactions with pathogens and herbivorous insects, especially in species with strong spatial variation and intense sib competition. These situations are plausible for trees with gravity-dispersed seeds (barochory). Additionally, separate from the fitness benefits to the parent tree, there exists (3) an effect of increasing the genetic variance of the whole population, which is important because the genetic variance determines the evolutionary ability of the population in responding to environmental changes.

### 5.2. Efficient elimination of deleterious mutations

An analysis of the next-generation sequencing data in trees concluded that somatic mutations behave as neutral variations during branch elongation but suffer deleterious effects when they form the next generations (Ally et al. 2010; Satake et al. 2023). This suggests that somatic mutations are maintained without receiving strong selection, which is quite different from the expectation of the standard population genetics theory predicting strong elimination of deleterious mutations (Otto & Orive 1995). This observation suggests a novel theoretical idea for the evolutionary advantage of tree tissue structure: trees might be able to effectively eliminate harmful deleterious genes, provided that the deleterious mutations have synergetic effects.

Many deleterious mutations of small effects occur in every generation and accumulate in the population without immediate elimination, reducing the population mean fitness (genetic load) and cause inbreeding depression. Some empirical evidence indicated that deleterious mutations work synergistically: the fitness of a genome including *k* deleterious genes is represented as *exp*[−*αk* − *βk*^2^] with *α* > 0 and *β* > 0 (Mukai 1969). This implies that the effect of a few mutations is small, but with a sufficiently high number of mutations, fitness declines rapidly in an accelerating manner (Maynard Smith 1978). One theory for the advantage of sexual reproduction concerns the efficient elimination of synergistically deleterious genes (Kondrashov 1988, 1993; Kondrashov & Crow 1988; Hurst & Peck 1996).

We can compare a tree that accumulate somatic mutations over hundreds of years with a large genet of a clonal plant that extends through many small individuals (ramets). In the clonal plant, deleterious mutations are expressed each time a ramet is formed, as shown in seagrass (Yu et al. 2020). In contrast, in tree, these mutations accumulate without being exposed to selection and only receive strong selection during the reproductive stages. Different branches (or different stem cells) of a tree may accumulate different number of deleterious mutations in a neutral manner. If these mutations interact synergistically during reproduction, a small fraction of seeds may carry many deleterious mutations, leading to significantly reduced fitness, while others may not exhibit as much deleteriousness. This, coupled with strong sib competition, could efficiently remove deleterious mutations and improve the reproductive success of the parent tree compared with the corresponding clonal plant.

At this moment, empirical evidence for the extent of synergism was not decisive: Synegism was supported in experiments with *Drosophila* (Mukai 1969) but not detected in a study with *E. coli* (Elena & Lenski 1997). Empirical studies on the accumulation of somatic mutations of trees and clonal plants yielded varying conclusions: Ally et al. (2010) reported a decline in fertility, Bobiwash et al. (2013) estimated the accumulation of deleterious mutations in a long-lived clonal shrub. In contrast, Cruzan et al. (2022) found within-individual selection in perennial herbs resulting in the production of advantageous mutations in some shoots, while Alejano et al. (2019) observed no correlation between fitness of offspring and the age of the parent tree. More empirical studies are needed.

### 5.3 Alternative indexes measure different aspects of novel genetic variation

In this paper, we discussed three indexes for the genetic diversity of somatic mutations within an individual tree. They quantify the amount of genetic diversity in different aspects.

The additive genetic variance of a population is crucial in determining its response to environmental changes. Somatic mutations in trees may increase the genetic variance of the population. To assess this effect, *D*_*PO*_ is important as it represents the magnitude of parent-offspring distance produced per generation.

In contrast, to measure the contribution to the fitness of the parental tree, we need to count the number of surviving offspring relative to other trees in the same population. Both 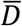 and *PD* measure the genetic diversity of offspring produced by each individual tree, rather than their effect to the entire population.

If we compare the two indexes, 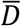 and *PD*, the phylogenetic distance *PD* is proportional to the total number of haplotypes included in the whole phylogeny. It assigns equal importance to genotypes with different number of seeds. In contrast, the mean phylogenetic distance between organs 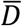 considers a branch with many seeds more important than another with only a few seeds.

In situations where the majority of seeds can survive and engage in intense sibling competition, the number of genetically distinct offspring becomes crucial, rendering multiple copies of the same genotype irrelevant. In such cases, *PD* (phylogenetic diversity) holds more significance than 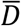 (mean pairwise distance). However, if a large fraction of offspring die early in their life before they start to compete with each other, a genotype existing as a single copy is likely to be eliminated stochastically. In such cases, producing many offspring of the same genotype makes sense to realize at least one individual can join in the sib competition. In such scenarios, 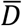, which considers the relative number of seeds, may be more appropriate than *PD*.

In a species-rich environment such as tropical rain forests, the specialist herbivores and pathogens are abundant near the parent tree (Janzen 1970). Having the offspring with traits different from the parent might be more important than having offspring that differ among themselves. In such a case, *D*_*PO*_ might be the most suited in evaluating the fitness contribution to the parent.

In short, the choice of the most suitable index for quantifying the genetic variation caused by somatic mutations should depend on which aspect of the many evolutionary effects one intended to evaluate.

To enhance our understanding of the role of tree tissue structure in accumulating somatic mutations, we require both theoretical studies to explore the evolutionary impacts of somatic mutations accumulated in trees and empirical research in the field that takes into account the success of all life stages, genetic structure, as well as both temporal and spatial variations.

## 7. Data and coding statement

Because the research is entirely theoretical, no empirical data is available. The datasets used and analyzed during the current study available from the corresponding author on reasonable request.

## 8. Acknowledgements

We thank the following people for their very helpful comments: R. Hayashi, R. Imai, S.P Otto, E. Sasaki, H. Tachida, T. Yahara, and R. Yamaguchi.

## 9. Financial support statement

This work was done in support of a Grant-in-Aid for Scientific Research for JSPS Fellows: 23KJ1722 to S.T. and Gants-in-aid for Scientific Research: 21H04781, 23H04965, 23H04966 to A.S. (Japan Society for the Promotion of Sciences)

## 10. Conflict of interest declaration

The authors declare that they have no conflict of interest.

## File S1: Supplemental Material

### Part A

#### Phylogenic diversity of cells within a single tree

Here, we explain how to evaluate the phylogenetic distance between two sampled cells and also how to assess the phylogenetic diversity among three or more sampled cells.

##### A.1 Phylogenetic distance

We denote the physical length along branch between the sampled cell X and the forking point fork[X, Y] as *L*(X, fork[X, Y]). Similarly we denote the length between cell Y and fork[X,Y] as *L*(Y, fork[X, Y]). The length of the path connecting X and Y along branches “after forking” is defined as the sum:

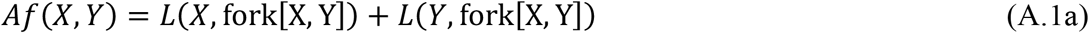

The phylogenetic distance between two sampled cells, X and Y, is

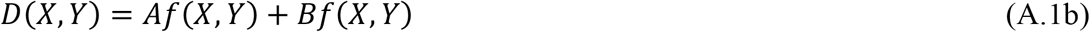

In a similar manner, we define the after-forking part of the distance between X and Z:

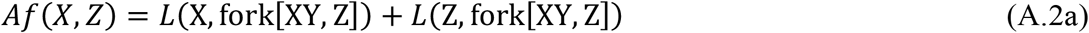

Note that fork[XY, Z] indicates that the location along shoot where the ancestral lineage of Z intersects with the common ancestral lineage of X and Y (See Fig. A1). The phylogenetic distance between X and Z is as follows:

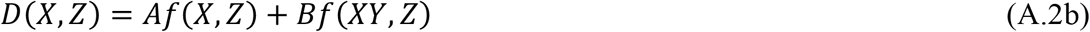

We can define the distance between Y and Z in a similar manner.

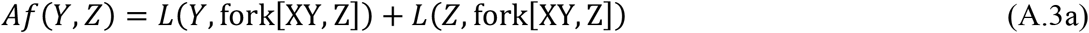

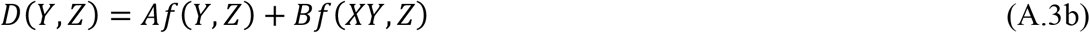

The before-forking portion is twice as long as the coalescent length discussed in Iwasa et al. (2023). It depends on the angle difference between two stem cells, indicating by *k* (*k* = 0, 1, 2, . . ., *n*). This corresponds to a radian angle of 2π*k/n*. It is challenging to determine the angle of the ancestral cells of a sampled cell located on a branch. Additionally, due to the failure of a stem cell to leave its successor, an ancestral stem cell lineage can occasionality be superseded by another stem cell line, causing a movement of the ancestral cell lineage (Iwasa et al. 2023). As a reesult, we randomize the angle difference and then consider the mean value of before-forking portion:

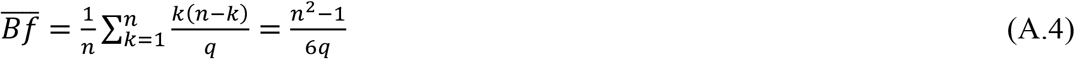

All the other measurements represent the physical distance along branches.

##### A.2 Phylogenetic diversity PD

The phylogenetic diversity of the sampled cells X, Y, and Z becomes as follows:

The undirected graph connecting X, Y, and Z by ancestral lineages is determined by summing the lengths of edges between the coalescent point between X and Y and the three sampled cells:

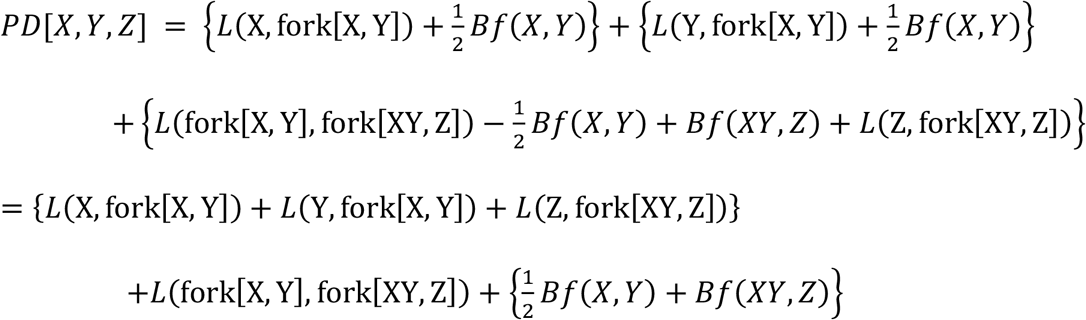

The last expression indicates that PD is the sum of terms calculated from the physical length along branches and the terms related to the before-forking portion. The former can be measured simply from the physical length, while the latter is connected tp the genetic diversity of stem cells within the SAM. This genetic diversity depends on the number of stem cells in the SAM, as well as the frequency of failure in asymmetric cell division and which stem cells supersede others when a stem cell fails to leave its successor stem cell (Iwasa et al. 2023).

### Part B

#### Derivation of the phylogenetic diversity (*PD*) of multiple sampled stem cells within the same SAM

Suppose we have sampled three cells at locations: *l*_0_, *l*_1_, and *l*_2_, which satisfy 0 ≤ *l*_0_ ≤ *l*_1_ ≤ *l*_2_ ≤ *n*. The intervals between these cells are denoted as *k*_1_, *k*_2_, *k*_3_, which are defined as follows: *k*_1_ = *l*_1_ − *l*_0_, *k*_2_ = *l*_2_ − *l*_1_, and *k*_+_ = *n* + *l*_0_ − *l*_2_. These intervals satisfy the following conditions:

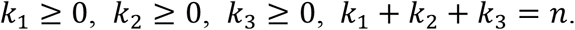

When one stem cell moves to the right by one, the interval in the left of the cell increases by one, which the interval in the right of the cell decreases by one. For example, when the *i*th cell moves to the right, *l*_*i*_ increases by one, *k*_*i*_ increases by one, and *k*_*i*+1_ decreases by one, simultaneously. If a cell moves to the left, the corresponding *l*_*i*_ decreases by one, *k*_*i*_ decreases by one, and *k*_*i*+1_ increases by one. The lengths of other intervals remain unchanged.

We first consider the case in which there are three cells, and *k*_1_ and *k*_2_ are integers within a triangular region on a plane: *k*_1_ > 0, *k*_2_ > 0, and *k*_1_ + *k*_2_ < *n*.

We adopt a continuous-time Markov chain, where only a single change occurs within a short interval. Let *D*(*k*_1_, *k*_2_, *k*_3_; *y*) represent the mean value of phylogenic diversity for three stem cells with angular intervals, *k*_1_, *k*_2_, *k*_3_ around the shoot axis, sampled at location *y* from the base of the shoot. Please note that, for the sake of brevity, we us the notation *D* in this appendix, which refer to *PD*. Then we derive the following recursive formula:

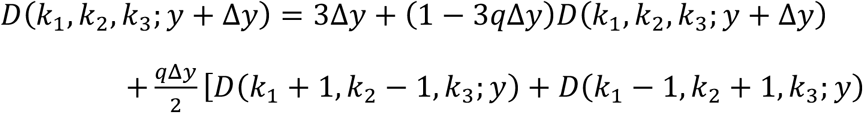

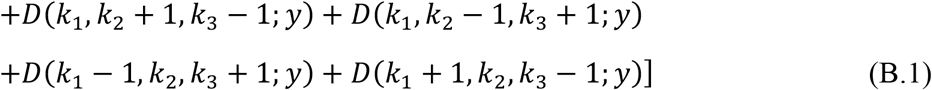

As Δ*y* approaches zero, we obtain

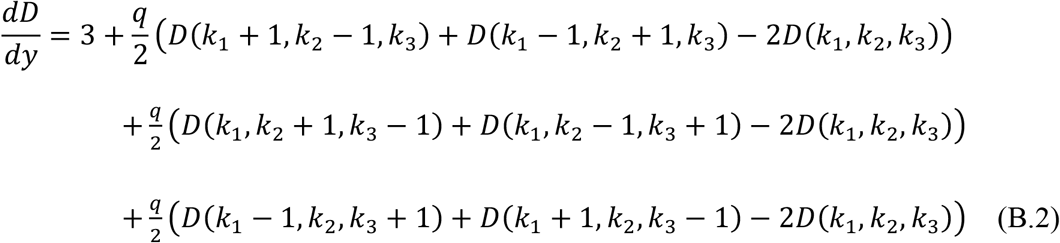

where (*k*_1_, *k*_2_, *k*_3_) represents all the points within the triangle. To save space, we do not display the dependence on *y*, although they are all functions that increase with *y*. In the limit of very large *y*, we have

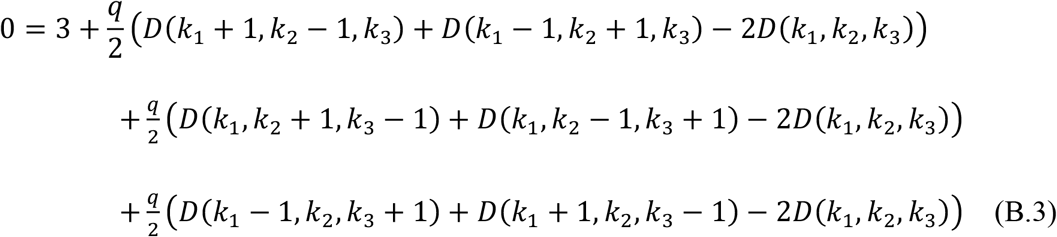

Here we introduce three operators, which modify a function *f*(*k*_1_, *k*_2_, *k*_3_). We define them as follows:

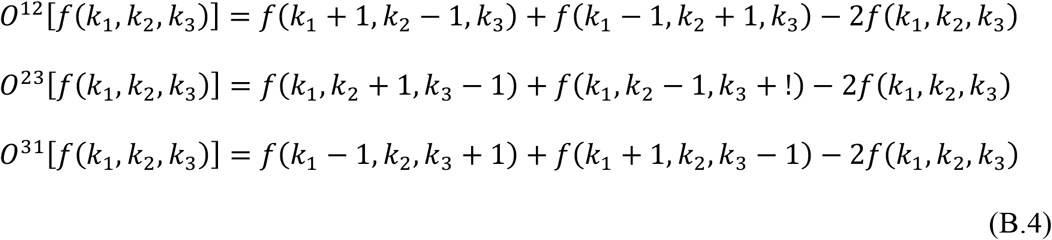

With these symbols, Eq. (B.3) is rewritten as follows:

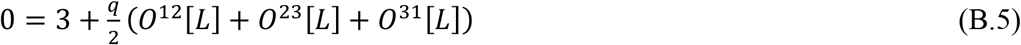

When we apply this to the function *f*(*k*_1_, *k*_2_, *k*_3_) = *k*_1_*k*_2_, we have the following results:

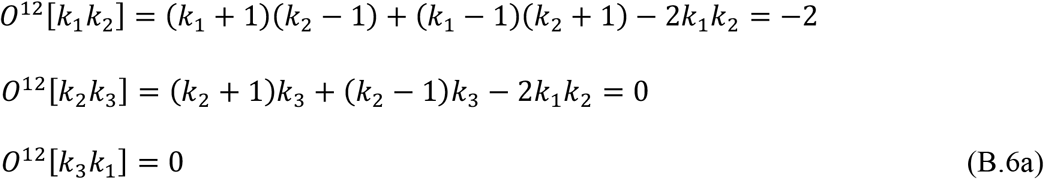

In a similar manner, we have

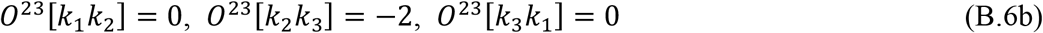

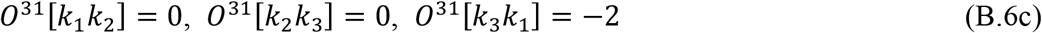

If we set *L*(*k*_1_, *k*_2_, *k*_3_) = *C*(*k*_1_*k*_2_ + *k*_2_*k*_3_ + *k*_+_*k*_1_), the following results holds:

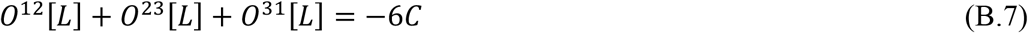

From Eq. (B.5), we have

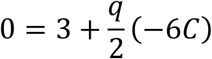

This implies that Eq. (B.3) holds if 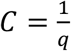. Therefore, one candidate solution for Eq. (B.3) is as follows:

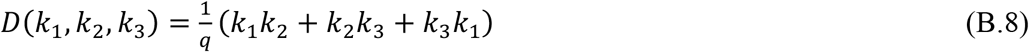

Hence, Eq. (B.8) satisfies Eq. (B.3). However, to be considered a valid solution, it must also meet the boundary condition. The boundary condition relates to the consistency of the solution when one of *k*_1_, *k*_2_, *k*_3_ is set to zero. In such a case, the coalescence of two of the three stem cells occurs, resulting in a model with two stem cells. This situation aligns with the model studied in Iwasa et al. (2023). Hence, we know that

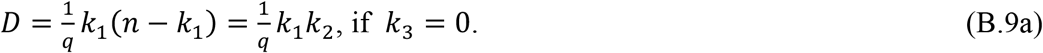

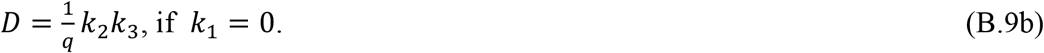

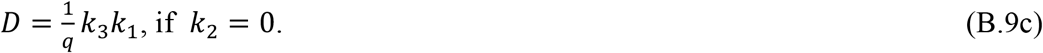

These conditions apply to the three edges of the triangular defined by Eq. (B.8). Consequently, we can conclude that *D*(*k*_1_, *k*_2_, *k*_3_) as presented in Eq. (B.8) is indeed the solution to (B.3). In other words, it is the solution to the differential equation (B.2) in the limit of as *y* approaches infinity.

Consider the scenario where a single coalescent event occurs. In this case, the operator of the differential equation is different from that of the three-stem cell case, as only two stem cells remain instead of three. In such a situation, the state point should lie on one of the three edges, rather than inside the triangle. Suppose it is the edge of *k*_3_ = 0. On this edge, the following equation must hold:

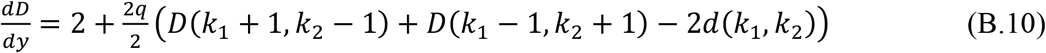

Note that the first term on the right-hand side is 2 instead of 3. This is because when one stem cell is replaced, both distances change, but we count the change in the distance twice because there are two stem cells involved. Both *k*_1_ and *k*_2_ must change if stem cells located at *l*_0_ or *l*_1_ are replaced. When *l*_1_ moves to the right by one step, *k*_1_ increases, and *k*_0_ decreases by one. If *l*_1_ moves to the left, the opposite changes occur. Additionally, the shift in *l*_0_ occurs at the same rate with similar effects, and hence we must multiply by a factor of 2. Therefore, we need to obtain the solution of the following:

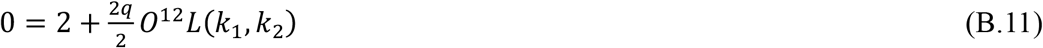

The solution of this equation is 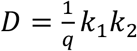.

If we consider four stem cells, the distance between adjacent stem cells are *k*_1_, *k*_2_, *k*_3_, and *k*_4_. The differential equation is as follows:

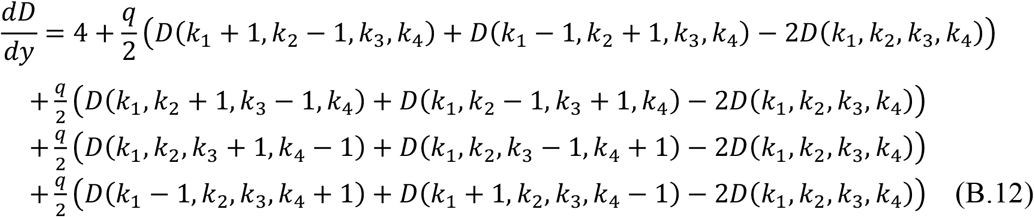

Building on this, we define operators in a manner similar to what was explained earlier, resulting in the following differential equation:

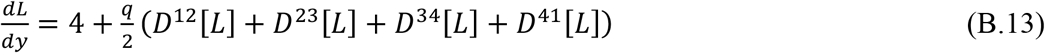

From this, the following represents a candidate solution for Eq. (B.13).

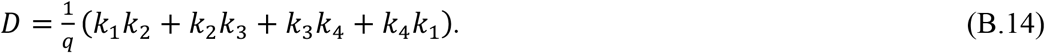

However, Eq. (B.14) does not satisfy the boundary condition on the plane *k*_4_ = 0. With *k*_4_ = 0, Eq. (B.14) yields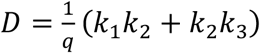, which is incorrect. The correct solution on the plane *k*_4_ = 0 is as follows:

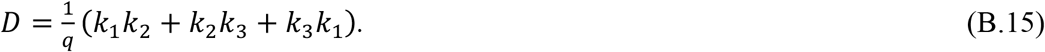

To obtain Eq. (B.15) on plane *k*_4_ = 0, we consider the following solution, which includes an additional term:

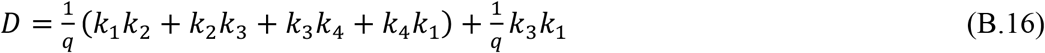

This additional term has no effect because it vanishes:

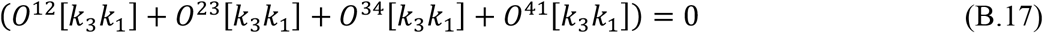

Therefore, *D* as defined by Eq. (B.16) is the same as Eq. (B.13) within the tetrahedron region.

Similarly, for consistency on planes *k*_1_ = 0, *k*_2_ = 0, and *k*_3_ = 0, we must introduce two additional terms:

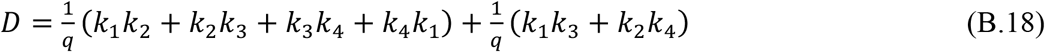

This aligns with Eq. (B.13) within the tetrahedron and is consistent with the boundary condition on the planes.

Based on these results, we propose that the following solution for the general case of *m*, the number of stem cells:

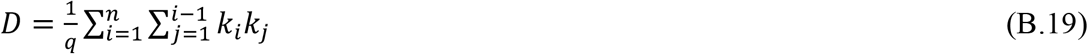

Note that the sum is calculated only once for each pair of stem cells. Because

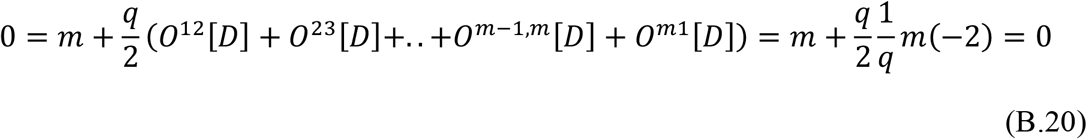

Therefore, Eq. (B.19) satisfies the differential equation.

The boundary condition is derived from the fact that when one of the intervals becomes zero, say *k*_*i*_ = 0, the system decreases in dimension by one. However, the operator must be appropriately adjusted.

Let us consider the following operator.

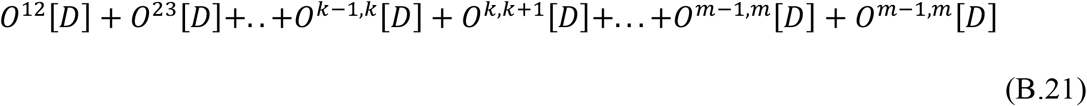

When *k* = 0, the one simply removed *O*^*k*−1,*k*^[*D*] and *O*^*k,k*+1^[*D*] is not appropriate:

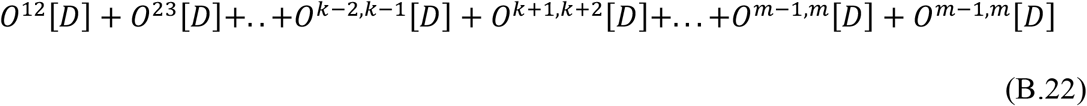

The correct one is the following:

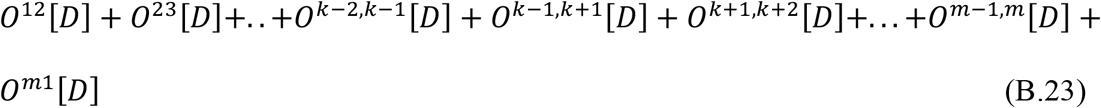

In this case, we need to introduce a new term *O*^*k*−1,*k*+1^[*D*]. Hence, when one interval disappears, a new term must be added.

Eq. (B.19) represents the total sum of the coalescent lengths of *m* stem cells. With this equation, we can rewrite *D* as follows:

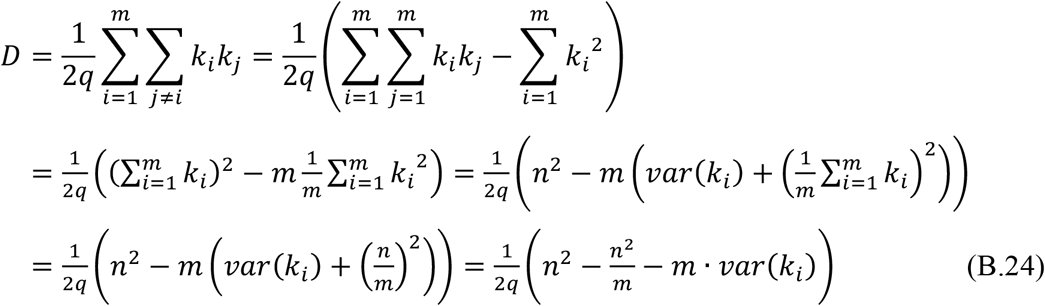

From this, we have

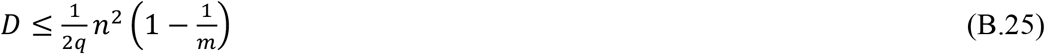

This quantity reaches the maximum when *m* stem cells are sampled at equal intervals. The equality holds, if *n* can be divided by *m* and the sampled cells are evenly distributed (*var*(*k*_*i*_) = 0).

When we sample only two cells (*m* = 2), we have 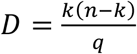 as derived in Iwasa et al. (2023). The maximum value of this is approximately 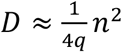, when sampled cells are positioned on the opposite side of the branch 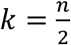.

When we sample all *n* stem cells in the shoot apical meristem, we obtain the total coalescent length by simply setting *m* = *n* and *k*_1_ = *k*_2_ =. . . = *k*_*m*_ = 1. Eq. (B.25) becomes as follows:

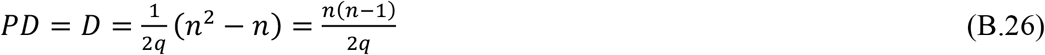

Here, we note that *D* is, in fact, phylogenetic diversity *PD* in the text. The ratio of the total coalescent length of the whole stem cells to the maximum coalescent length when only two cells are sampled is

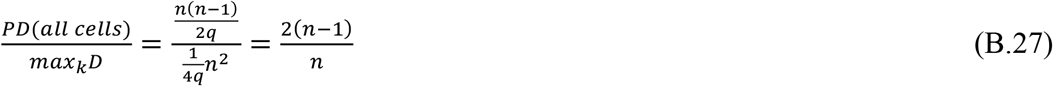

which is slightly less than 2.

We can consider the value of *D* averaged over *k* (*k* = 1,2, . . ., *m*). This is the mean pairwise phylogenetic distance, denoted as 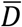 in the main text. It is calculated as follows:

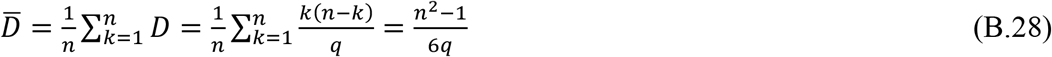

Refer to Iwasa et al. (2023). The ratio of phylogenetic diversity (the total coalescent length of all stem cells) to pairwise phylogenetic distance (the maximum coalescent length when only two cells are sampled is

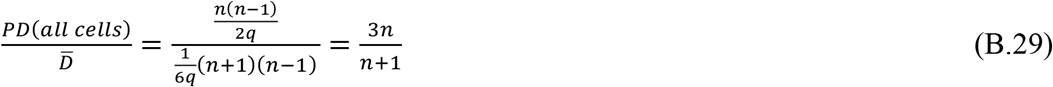

which is slightly less than 3.

In summary, the phylogenetic distance of all stem cells in the SAM is approximately three times larger than the mean pairwise phylogenetic distance and about two times larger than the maximum phylogenetic distance between two stem cells.

### Part C

#### Selective effects of genetic diversity in offspring

The genetic variation of phenotypic traits is an indispensable element of evolutionary adaptation. However, mutation also produces maladaptive or nonfunctional genes. We may ask whether having a higher rate of mutation is advantageous in the evolution and whether the tissue structure favoring somatic mutations could be selected for because of its contribution to enhanced genetic diversity. In the following subsection, we briefly summarize major theoretical ideas on the evolution of mutation rate in the population genetics.

i. Constant and uniform environment Theoretical study of the evolution of mutation rate includes papers on the evolution of genetic systems (Karlin 1975; Karlin & McGregor 1974). In the simplest setting of the model, there is one selected locus that affects fitness and a second modifier locus that specifies the mutation rate at the selected locus. The modifier may not directly contribute to fitness (a neutral modifier), but it evolves through its impact on the selected locus. In an infinitely large population of haploid organisms, if the allele fitness does not change over time, natural selection causes the first locus to converge to the state of the highest fitness. The population comes to have alleles at a local peak of the fitness landscape. Then the mutations become harmful because they change the optimal allele to a different allele with lower fitness. Hence, the mutation rate evolves to zero. This argument was extended to various situations with nonlinear fitness effects, such as diploid with dominance between alleles, multiple loci with epistasis, and phenotypic switching (Liberman et al. 2011; Altenberg et al. 2017).
ii. Temporally fluctuating environments Under temporally changing environments, alleles advantageous in the current generation may become disadvantageous, while those that will be advantageous later are currently disfavored and rare. The mutation creates promising alleles from those that are currently favored and abundant. A positive mutation rate evolves in a cyclical environment (Ishii et al. 1989), but not in randomly changing environments. This result can be intuitively understood by decomposing the environmental fluctuation into a sum of multiple sinusoidal components with varying periods, provided that selection is not very strong. In an environment fluctuating with a single sinusoidal component, the mutation rate to evolve is inversely proportional to the period. Hence, a high mutation rate evolves under fluctuation with a short period, while a low mutation rate evolves under fluctuation of a long period. On the other hand, the impact of environmental fluctuations on the mutation rate modifier gene and the population mean fitness is weaker for components with short periods (fast fluctuations) than for those with long periods (slow fluctuations). This phenomenon is known as “low-pass filter effect,” reported earlier in the theory of recombination rate evolution under fluctuating environments (Sasaki & Iwasa 1987). Furthermore, random fluctuations contain long-period components more than short-period components, as indicated by the monotonic decrease in the power spectrum with the frequency. Consequently, random fluctuations do not make a positive mutation rate evolve, unless selection intensity is very strong. However, high mutation rates can evolve under frequency-dependent selection. In the host-pathogen and herbivore-plant coevolution with genotype-specific attacking rates, the fitness is likely to be frequency dependent and a high mutation rate evolves (Haraguchi & Sasaki 1996, 1997; Sasaki & Haraguchi 2000).
iii. Spatial heterogeneity and severe sib competition. Another mechanism promoting the evolution of a positive mutation rate considers sib competition. This logic was originally proposed as a mechanism for the evolution of sexual reproduction with recombination (Williams 1975), and was later formalized in a mathematical model (Maynard Smith 1978; Douge & Iwasa 2017a, 2017b), where an element of spatial heterogeneity was incorporated, as a concept known as the “tangled bank” hypothesis (Bell 1982). Suppose that the habitat consists of many patches and that the environmental condition changes between patches and between generations. Even if a parent’s allele is well adapted in the current generation, it may not be adapted in the next generation. If competition is very intense, the best-adapted individuals among the seeds that landed on the same patch take over the patch. The advantage of producing many offspring of the same genotype is much reduced if multiple offspring of the same parent land on the same patch and engage in severe sib competition. In such a situation, a parent with a high mutation rate receives an advantage by producing genetically more diverse offspring. This scenario seems plausible for many trees that have a limited range of seed dispersal and produce a large number of seeds, leading to intense sib competition. Many trees that rely on gravity-dispersal, such as those producing acorns, have a limited seed dispersal range. However, the effect might be weaker for trees producing seeds with a long dispersal distance, such as those carried by ants or other animals using elaiosomes, as well as small seeded species like maples and other gap adapted species.
iv. Mutation enhancing mechanisms In special occasions, organisms adopt high mutation rates (Metzgar & Willis 2000). In bacteria, a high mutation rate is induced in response to risky environmental conditions, as exhibited by the SOS mechanism (Rosenberg et al. 1998). Other organisms, such as parasite worms like trypanosomes, have genes for surface proteins that allow them to escape host immune responses.

From these observations, we must conclude that a higher rate of mutations in general is disadvantageous, except for specific mechanisms for coping with host-pathogen interaction. An observed positive mutation rate is a result of a large cost accompanied by further reductions in error rates during genome replication and damage repair. A small positive rate of mutation produces genetic variation that forms the basis for the adaptation to environmental changes (Lynch 2010, 2011).

## Notes

### Competing Interest Statement

The authors have declared no competing interest.

